# On the critical concentration for net assembly of dynamically unstable polymers

**DOI:** 10.1101/2024.04.12.589322

**Authors:** R. Dyche Mullins

## Abstract

Cytoskeletal and cytomotive filaments are protein polymers that move molecular cargo and organize cellular contents in all domains of life. A key parameter describing the self-assembly of many of these polymers —including actin filaments and microtubules— is the minimum concentration required for polymer formation. This ‘critical concentration for net assembly’ (cc_N_) is easy to calculate for eukaryotic actins but more difficult for dynamically unstable filaments such as microtubules and some bacterial polymers. To better understand how cells (especially bacteria) regulate assembly of dynamically unstable polymers I investigate the microscopic parameters that influence their critical concentrations. Assuming simple models for spontaneous nucleation and catastrophe I derive expressions for the monomer-polymer balance. In the absence of concentration-dependent rescue, fixed catastrophe rates do not produce clear critical concentrations. In contrast, simple ATP-/GTP-cap models with concentration-dependent catastrophe rates, generate phenomenological critical concentrations that increase linearly with the rate of nucleotide hydrolysis and decrease logarithmically with the rate of spontaneous nucleation.

## Introduction

Many cellular processes require assembly and/or disassembly of cytoskeletal structures — networks of protein polymers that generate, resist, and transmit forces over cellular length scales. These polymers include eukaryotic microtubules and actin filaments, along with their bacterial and archaeal relatives (Brown, 2023; Charles-Orszag, 2024).

Actin filaments assemble via a nucleation-condensation mechanism, and exhibit a *critical concentration* for net polymer assembly. Kasai and co-workers (1962a) first defined this critical concentration as the concentration above which the amount of filamentous actin increases linearly with increasing total actin concentration. These authors also noted that this value represents the concentration of actin monomers in equilibrium with polymer. Net assembly of microtubules also occurs above a definite critical concentration (Gaskin, 1974; Johnson, 1975), but the biophysical origin of *this* critical concentrations turns out to be different from that of actin. The reason for this difference is that actin filaments undergo *reversible polymerization* while microtubules exhibit *dynamic instability*.

Holy and Leibler define reversible polymerization as “a continuous competition between […] assembly and disassembly” (Holy, 1994). Under these conditions, the critical concentration for net polymer assembly is the concentration at which the rates of subunit addition and loss are balanced (Kasai, 1962a; Kasai, 1962b). At steady state these filaments do not grow; instead their lengths exhibit a relatively slow, one-dimensional random walk as subunits bind and dissociate with equal frequencies (Holy, 1994).

In contrast, dynamically unstable polymers switch stochastically between reversible polymerization and rapid, catastrophic shortening (Mitchison, 1984; Horio, 1986). As a consequence, the critical concentration for net assembly of these polymers is not determined solely by the rates of elongation and shortening, but also depends on the balance between filament nucleation and catastrophic filament loss. This discrepancy was first noted by Mitchison and Kirschner (1984) and described more clearly by Walker et al. (1988), who defined two critical concentrations that govern microtubule dynamics: one for net assembly and one for filament elongation (cc_N_ and cc_E_ in my nomenclature). The same two critical concentrations have been observed experimentally in dynamically unstable actin-like filaments from bacteria (Garner, 2004; Petek, 2017).

The critical concentration for net assembly will always be higher than that for elongation (cc_N_ > cc_E_) but, in principle, the two values can be arbitrarily close. Experimentally, however, values of cc_N_ measured for all dynamically unstable polymers to date turn out to be significantly higher than those for cc_E_ (Walker, 1988; Garner, 2004; Petek, 2017). This relatively large mismatch between cc_N_ and cc_E_ enables dynamically unstable filaments to maintain a monomer concentration high enough to support filament elongation at steady state. This is important in bacteria, where assembly of actin-like polymers, such as ParM, can move cellular cargo under steady-state conditions (Garner, 2007; Campbell, 2007), without relying on the disassembly factors and recycling proteins of the type that maintain the steady-state actin monomer concentration in eukaryotic cytoplasm well above cc_E_ (Loisel, 1999; Pollard, 2000).

Dogterom and Leibler (1993) made a seminal contribution to the study of dynamic instability by calculating a critical tubulin concentration that marks a transition from bounded to unbounded microtubule growth. These authors proposed a simple, two-state model in which microtubules grow from fixed nucleation sites and switch stochastically from growing to shortening (catastrophe) and back again (rescue). By assuming (i) a single-step catastrophe mechanism with a fixed frequency and (ii) a rescue frequency that increases linearly with monomer concentration, Dogterom and Leibler identified a soluble tubulin concentration at which rescue compensates catastrophe and filaments begin to grow without bound. In a limiting pool of total tubulin, this critical concentration becomes equivalent to cc_N_.

The transition from bounded to unbounded microtubule growth described by Dogterom and Leibler (1993) depends absolutely on the assumption of frequent and monomer-dependent rescue. For this reason the model cannot be invoked to calculate or even satisfactorily explain cc_N_ in bacterial actin-like polymers, such as ParM or Alp7A, whose depolymerizing filaments exhibit little or no evidence of spontaneous rescue (Garner, 2004; Garner, 2007; Rivera, 2011). To understand the molecular mechanisms behind rescue-independent critical concentrations, I derive general expressions for monomer-polymer balance of a dynamically unstable filament, assuming spontaneous, asynchronous, and concentration-dependent nucleation and no independent rescue process. These assumptions better reflect the behavior of bacterial actin-like proteins such as ParM and Alp7A and the conditions under which they function in the cell. I then investigate the consequences of two simple models for catastrophe: (i) a fixed rate mechanism and (ii) stabilization by a simple ATP-/GTP-cap. Interestingly, in the absence of monomer-dependent rescue, fixed catastrophe rates produce nonlinear polymer accumulation but no clear critical concentration. In contrast, nucleotide-cap models, which predict concentration-dependent catastrophe, produce sharper polymer transitions and a phenomenological critical concentration. This critical concentration increases linearly with rate of nucleotide hydrolysis; decreases logarithmically with the rate spontaneous nucleation; and helps explain steady-state assembly of filament structures in bacterial cytoplasm.

## Results and Discussion

Here I consider polymers that switch stochastically between reversible polymerization and catastrophic shortening, with no mechanism for switching back to stable growth once catastrophe has occurred. Assuming that new filaments appear by spatially distributed, asynchronous, and spontaneous nucleation, the steady-state number concentration of *filaments* is given by the product of the rate of nucleation (*R*_*N*_) and the average filament lifetime (<*t*_*L*_>). Multiplying this product (*R*_*N*_<*t*_*L*_>) by the population average of filament lengths, <*L*_*av*_> (subscript *av* denotes mean length over a filament’s lifetime; pointed brackets denote a population average of these mean lengths), we obtain an estimate of the polymer concentration. If lengths and lifetimes of individual filaments were uncorrelated this estimate would be accurate. Longer filaments, however, take longer to assemble and disassemble and so lifetime and length are strongly correlated. An accurate value for polymer concentration ([*p*]) requires calculating the population average of the *product*: *t*_*L*_*L*_*av*_ (**Eqn. 1**).

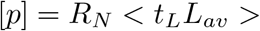

In this equation length is expressed in number of protomers, and thus unitless.

### A general expression for polymer concentration

Along with spontaneous nucleation I now make three additional assumptions: (1) filaments grow at a constant rate determined by the soluble monomer concentration and rate constants for elongation and shortening (*k*_*+*_ and *k*_*-*_); (2) elongation is terminated by major catastrophes that occur with some characteristic frequency distribution; and (3) catastrophic shortening always leads to filament loss (i.e. no “rescue”). The first two assumptions are generally valid. The third assumption holds for bacterial actin-like filaments ParM and Alp7A, but does not always apply to eukaryotic microtubules. For a given filament growing and shrinking under these conditions a plot of length versus time describes a triangle whose base is the filament lifetime and whose height is its maximum length **(Figure 1)**. When growth and shortening are both linear, the average length of a filament over its lifetime is half its maximum length. The product of lifetime and average length, therefore, is equivalent to the area under the triangle (*t*_*L*_*×L*_*max*_*/2*). The length of the base of this triangle is related to its height by the slopes of steady elongation (*s1*) and catastrophic shortening (*s2*).

**Figure 1.**
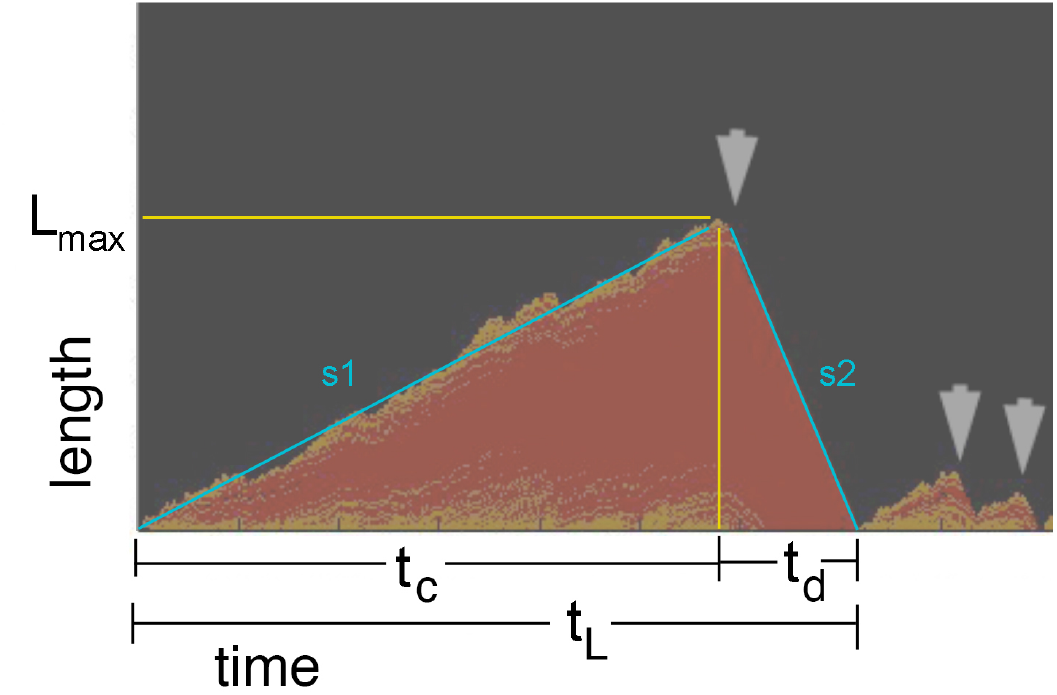
Calculating the product of filament length and lifetime. The plot represents growth and shortening of a dynamically unstable filament. YELLOW: ATP-bound protomers forming caps; RED:ADP-bound protomers. L_max_: maximum filament length. T_c_ and t_d_: times of assembly and disassembly phases. t_L_: total filament lifetime. Slopes, s1 and s2, are rates of elongation and catastrophic shortening.

The total filament lifetime is the sum of the time spent elongating to catastrophe (*t*_*c*_) plus the time spent depolymerizing to extinction (*t*_*d*_). The first term is the maximum length divided by the rate of filament elongation.

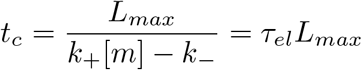

And the second term is the maximum length divided by the rate of catastrophic shortening

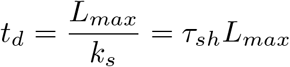

So the total lifetime (*t*_*L*_) of a filament whose length is limited by major catastrophes is

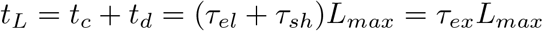

We define *τ*_*el*_ and *τ*_*sh*_ as the characteristic elongation and shortening times and the sum, *τ*_*ex*_, as the characteristic *excursion* time associated with the addition and loss of a single subunit from the filament. The product of filament lifetime and time-averaged length is, therefore, given by

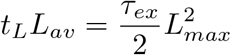

From this equation we can see that the population average of the product of filament length and lifetime is proportional to the *second moment* of the length distribution.

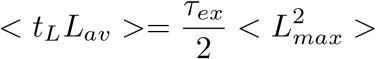

For filaments growing at a constant rate, *t*_*c*_ and *L*_*max*_ are related by the characteristic elongation time, *τ*_*el*_. Calculating moments of the *maximum length* distribution is, therefore, equivalent to calculating moments of the *catastrophe time* distribution (**Eqn. 2)**.

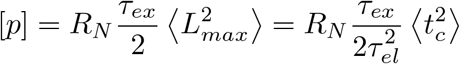

Note that **Eqn. 2** will hold for a variety of catastrophe time distributions —e.g. for both single- and multi-step catastrophe mechanisms

### Polymer concentration for single-step (and approximately single-step) catastrophe mechanisms

We will now add the assumption that catastrophe times are exponentially distributed, a case that corresponds to a single-step mechanism for catastrophe initiation. This assumption is likely not valid for eukaryotic microtubules (Gardner, 2013), but does apply to a broad range of ATP-/GTP-cap models of dynamic instability, including those that likely govern bacterial actin-like filaments, such as Alp7A and ParM. For this reason, the remainder of my analysis and the example cases considered will focus on dynamically unstable bacterial actin-like proteins.

For a single-step catastrophe mechanism, both the catastrophe times and filament lengths will be exponentially distributed, and the probability density function describing catastrophe times is given by

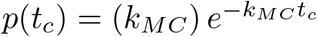

Where *k*_*MC*_ is a rate parameter for *major catastrophes*—switching events that lead to complete extinction of the filament. These events are also called *global catastrophes* (e.g. by Antal, 2007a and 200b). The second moment of this distribution is

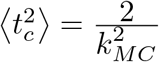

Substituting this value back into **Eqn. 2** yields (**Eqn. 3**):

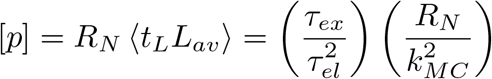

This expression makes only one assumption about catastrophe, namely that catastrophe times are exponentially distributed. While this assumptions holds *exactly* for single-step mechanisms, it turns out to be *approximately* true for a large class of more complicated ATP/GTP-cap models (Dieterle, 2022), making this equation more general than initially advertised.

### Incorporating simple models for nucleation and catastrophe

For actin and actin-like polymers the nucleation rate scales approximately as the third power of the free monomer concentration (Kasai, 1962b; Garner, 2004; Petek, 2017).

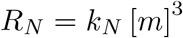

Where *k*_*N*_ is a constant. If, like Dogterom and Leibler, we assume a constant catastrophe frequency —independent of elongation rate— then polymer concentration is given by

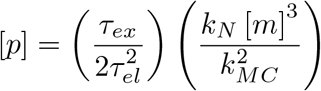

Here, monomer-dependent nucleation balances a constant catastrophe rate, but does not produce a clear critical concentration (cc_N_). This can most easily be seen in the special case of irreversible elongation (*k*_*-*_=0), where cc_E_ is zero. If we further assume fast catastrophic disassembly (*τ*_*sh*_=0), then *τ*_*el*_ = *τ*_*ex*_ = 1/*k*_*+*_[*m*] and polymer concentration becomes (**Eqn. 4**)

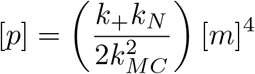

This function generates no sharp, monomer-dependent increase in polymer concentration (Figure 2a). Additionally, the concentration of monomer in equilibrium with polymer does not remain constant but increases as the fourth root of total protein concentration (Figure 2b).

**Figure 2.**
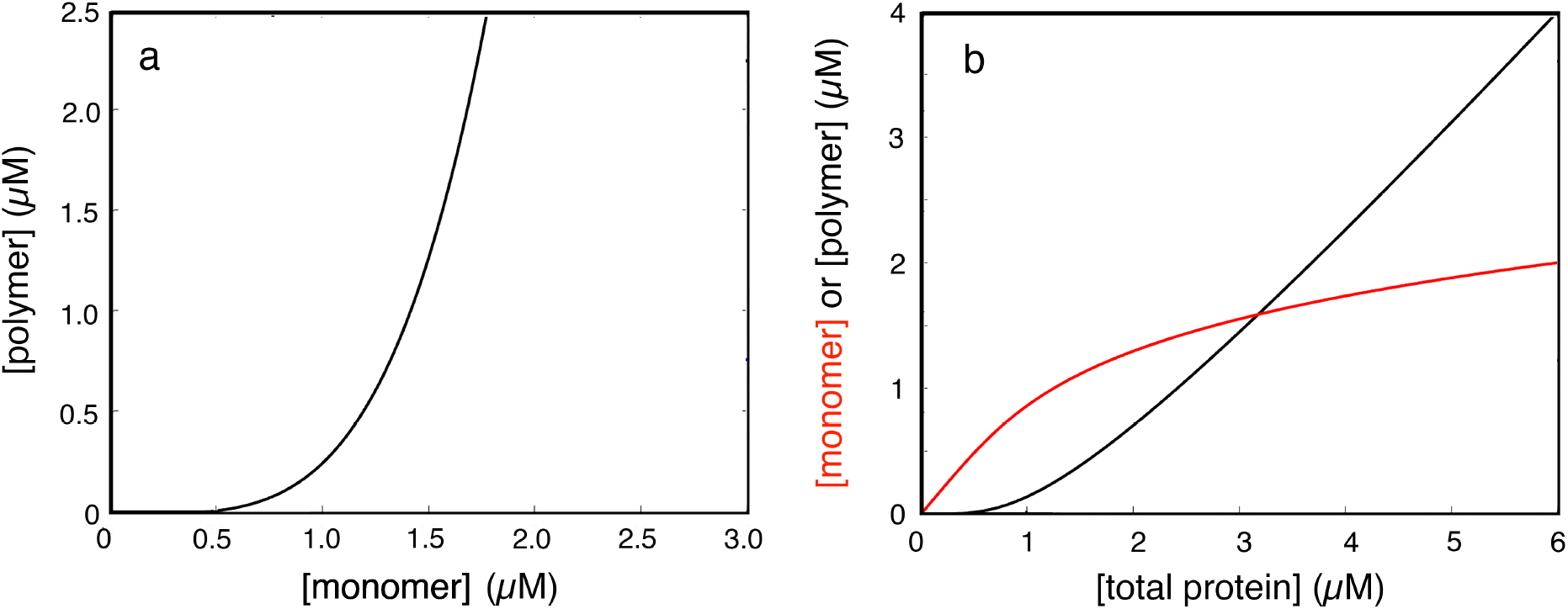
Monomer-polymer relationships for a fixed catastrophe rate (*k*_*MC*_ = 0.01), according to **Eqn. 4**. (a) Polymer concentration as a function monomer for physiologically relevant values of biochemical parameters (Table 1). (b) Monomer and polymer concentrations as a function of total protein. The plot was generated by numerically inverting **Eqn. 4** using parameter values as in (a).

### Nucleotide cap models for catastrophe

I next consider models in which stable elongation depends on the presence of ATP- or GTP-bound protomers at the filament end (Mitchison, 1984). In the simplest case, where one ATP-bound protomer suffices to stabilize the filament, the catastrophe rate (*k*_*MC*_) is given by an infinite product with an elegant asymptotic expansion (Apostol, 1976; McIntosh, 1998):

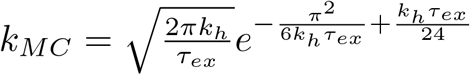

Related expressions can be derived for more complicated cap models and this will be the subject of a future paper (also, see below). For now, substituting expressions for nucleation and catastrophe into **Eqn. 3** yields a closed-form expression for polymer concentration as a function of soluble monomer (**Eqn. 5**):

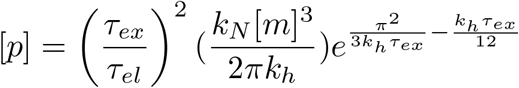

As above, consider the limiting case of irreversible polymerization (*k*_*-*_ = 0) and fast catastrophic shortening (τ_sh_ = 0). The characteristic elongation and excursion times (*τ*_*el*_ and *τ*_*ex*_) both reduce to 1/*k*_*+*_[*m*], and the expression for polymer concentration becomes (**Eqn. 6**)

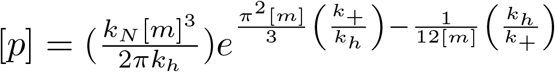

The [*m*]^*3*^ and especially the exponential terms in **Eqn. 6** make the polymer concentration very nonlinear with respect to the free monomer concentration (**Figure 3a**). At very low monomer concentrations polymer is almost non-existent. Between 0.2-0.4 µM monomer, however, polymer concentration explodes, increasing by more than four orders of magnitude (**Figure 3a**). By the time monomer concentration reaches 2 µM polymer concentration has risen to ∼10^30^ µM. Mathematically, both the suppression and the rapid growth of polymer are driven by the exponential term in **Eqn. 6**. At low monomer concentrations the exponential term is dominated by *e*^(-*k*_*h*_/12*k*_*+*_[*m*]), which drives the polymer concentration down near zero. Conversely, at high monomer concentrations the exponential term is driven by *e*^(*π*^2^*k*_*+*_[*m*]/3*k*_*h*_), which rapidly pushes the polymer concentration to extremely high values (**Figure 3a**, inset). The transition point where the exponential term in **Eqn 6** shifts from suppressing polymer concentration to driving explosive growth occurs in the region where the exponent equals zero, i.e.

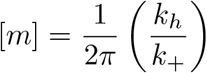

This transition resembles the shift from bounded to unbounded microtubule growth described by Dogterom and Leibler (1993). Instead of reflecting the balance of catastrophe and rescue, as in Dogterom (1993), however, this behavior arises from the effect of elongation rate on ATP-/GTP-cap size and catastrophe frequency.

**Figure 3.**
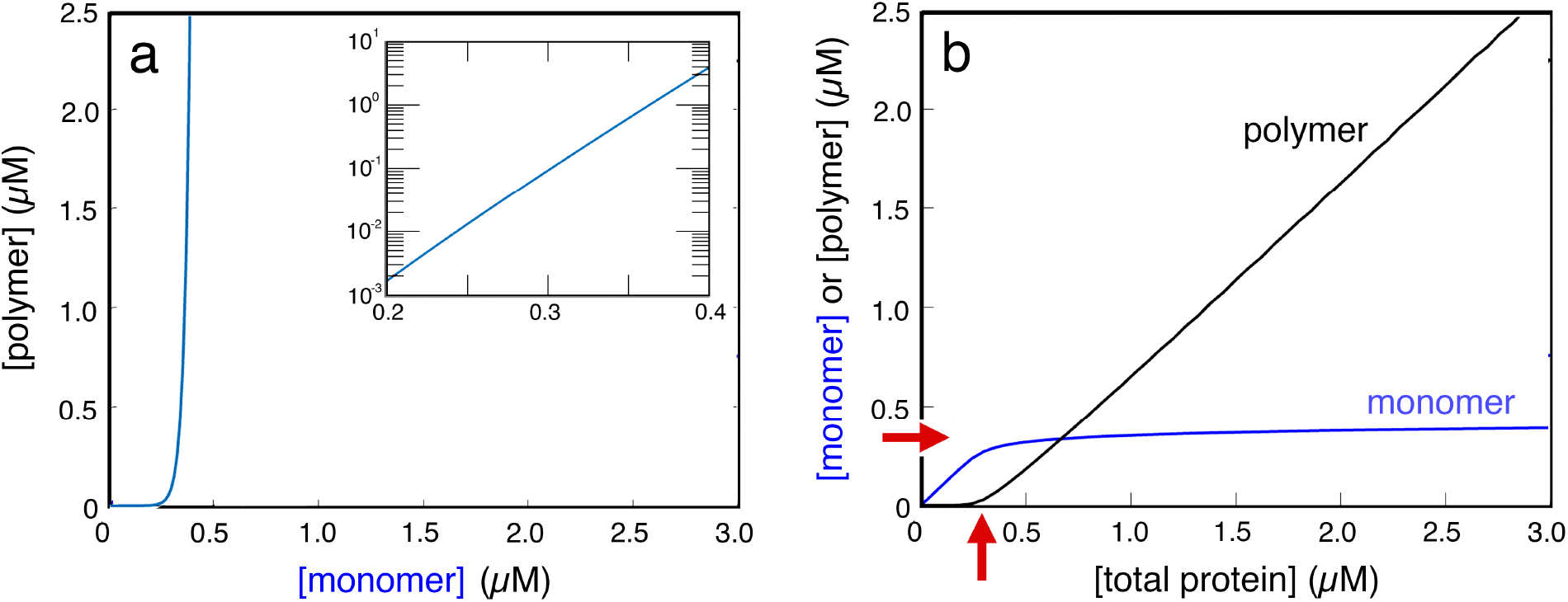
Relationships between concentrations of monomer, polymer, and total protein for a dynamically unstable filament according to **Eqn. 6**. (a) Polymer concentration as a function monomer for physiologically relevant values of biochemical parameters (Table 1). Note the the steep takeoff in polymer concentration around 0.3 µM monomer. Inset: log-log plot of polymer concentration in the takeoff region (0.2-0.4 µM monomer). In this window the polymer concentration increases ∼4 orders of magnitude. (b) Plot of monomer and polymer concentrations as a function of total protein. The plot was generated by numerically inverting **Eqn. 6** using parameter values as in (a). The steep take-off observed in (a) ensures that, beyond a concentration of ∼0.3 µM, the monomer concentration grows very slowly with total protein (∼[total]^1/30^). Red arrows mark the steady-state monomer concentration (on vertical axis) and critical-concentration for net polymer assembly (on horizontal axis).

**Table 1.** Values used (except as noted) in Figures 2-6.

|  |  |  |
| --- | --- | --- |
| Based on Petek (2017) and Garner (2004) |  |  |
| $k_+$ | $5 \mu M^{-1} sec^{-1}$ | ATP-monomer elongation |
| $k_-$ | $2 sec^{-1}$ | ATP-monomer dissociation |
| $k_-/k_+$ | $0.4 \mu M$ | Elongation crit. conc. ($cc_E$) |
| $k_s$ | $67 sec^{-1}$ | ADP-monomer dissociation |
| $k_h$ | $0.5 sec^{-1}$ | ATP hydrolysis |
| $k_N$ | $0.001 \mu M^{-2} sec^{-1}$ | nucleation |

Solving **Eqn. 6** numerically and plotting both monomer and polymer as a function of total protein concentration (**Figure 3b**), we see that the monomer concentration rises quickly at first and then approaches an apparent plateau as more protein is added. In the first phase, almost no polymer is present but then, abruptly, polymer begins to accumulate almost linearly while the monomer concentration remains almost constant. This apparent plateau in monomer concentration is not an asymptote, but because of the highly nonlinear character of **Eqn. 6**, monomer increases extremely slowly —as the ∼1/30^th^ power of total protein concentration. This behavior is phenomenologically similar to that of actin near the critical concentration for net assembly (Kasai, 1962) —i.e. polymer begins to increase linearly, but only after total protein concentration rises above a non-zero threshold value (**Figure 3b**, vertical arrow). Note that this threshold value approximately equals the monomer concentration in equilibrium with polymer (**Figure 3b**, horizontal arrow). This remains true across the physiologically relevant range of protein concentrations (<50µM). We, therefore, refer to this threshold concentration as a *phenomenological* critical concentration for net assembly.

We can approximate the critical concentration for net assembly as the point where the monomer and polymer concentrations are equal (**Figure 3b**), which enables us to transform **Eqn. 6** into the following expression (**Eqn. 7**)

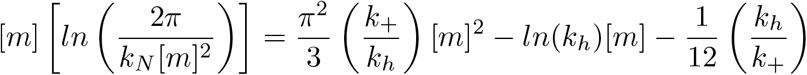

This equation separates the rate constants governing nucleation and catastrophe to the left and right sides respectively. We can solve this equation graphically by plotting the two sides separately and finding the point where they intersect (**Figure 4**). The right side of **Eqn. 7** is a quadratic function of [*m*] with coefficients that depend on rate constants for elongation and nucleotide hydrolysis. The left side of **Eqn. 7** is approximately linear in [*m*] with a slope that depends on the rate constant for nucleation.

**Figure 4.**
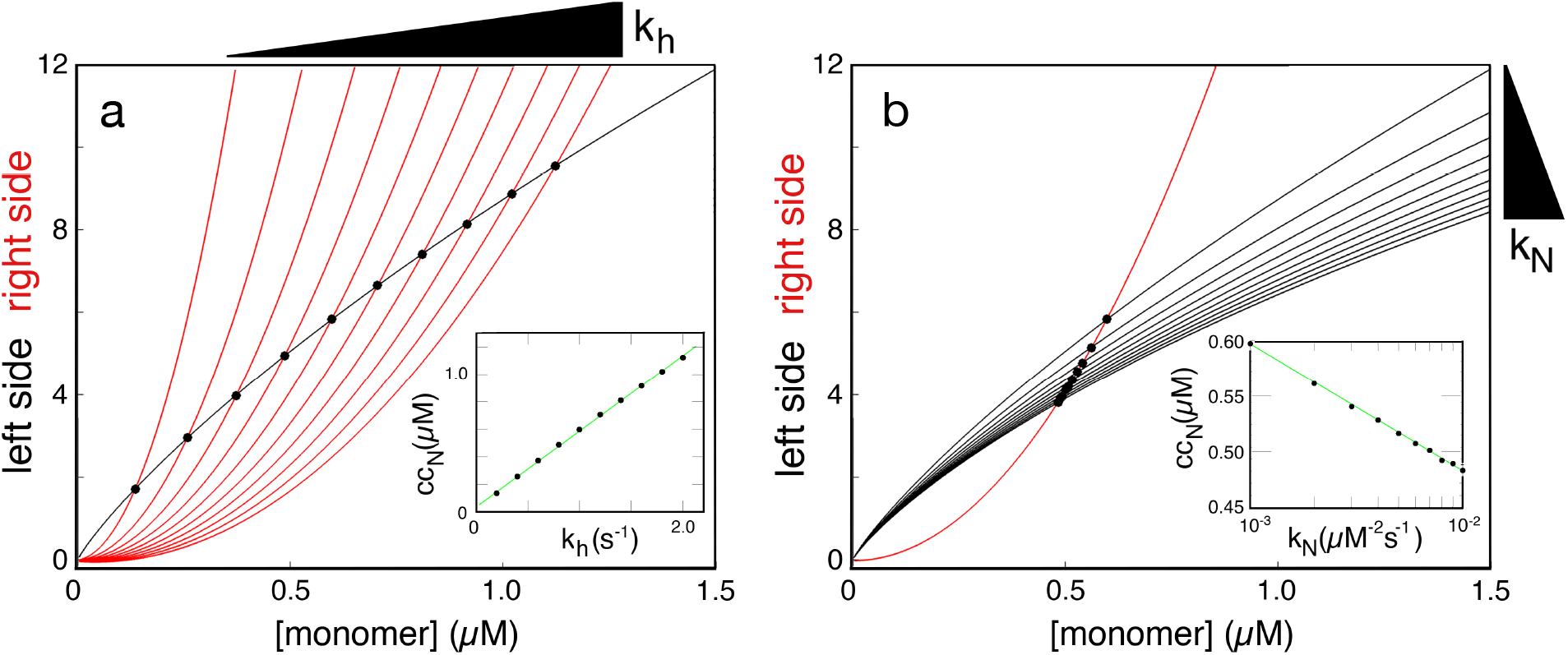
Graphical solutions to **Eqn. 7** for varying values of kh (a) and kN (b). In both panels the left-hand side of **Eqn. 7** is plotted in black; the right hand side is plotted in red. The intersection of the two curves —marked with a black circle— represents the critical concentration. In (a) *k*_*N*_ =0.001 µM-2s-1, *k*_*+*_ =5 µM-1s-1 and *k*_*h*_ has the values: 0,2, 0.4, 0.6, 0.8, 1.0, 1.2, 1.4, 1.6, 1.8, and 2.0 s-1. The black triangle above the red curves denotes the direction of increasing *k*_*h*_. Inset: computed critical concentrations as a function of nucleotide hydrolysis rate. In (b) *k*_*h*_=1s-1, and *k*_*N*_ has the values: 0.001, 0.002, 0.003, 0.004, 0.005, 0.006, 0.007, 0.008, 0.009, 0.01 µM-2s-1. The black triangle to the right of the black curves denotes the direction of increasing *k*_*N*_. Inset: computed critical concentrations as a function of nucleation rate constant —x-axis is logarithmic.

Varying the rate constants for nucleotide hydrolysis (**Figure 4a**) and filament nucleation (**Figure 4b**) in the physiologically relevant range for bacterial actin-like proteins (Table 1) reveals that

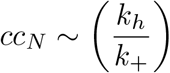

and

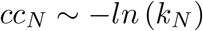

Now consider the more general case of reversible polymerization (k-> 0) and finite shortening rate (τsh > 0). Using **Eqn. 5** to calculate cc_N_ with parameter values from Table 1 and and a range of values for *k*_*h*_ and *k*_*N*_ (**Figure 5**) reveals similar scaling relationships to those in **Figure 4**.

**Figure 5.**
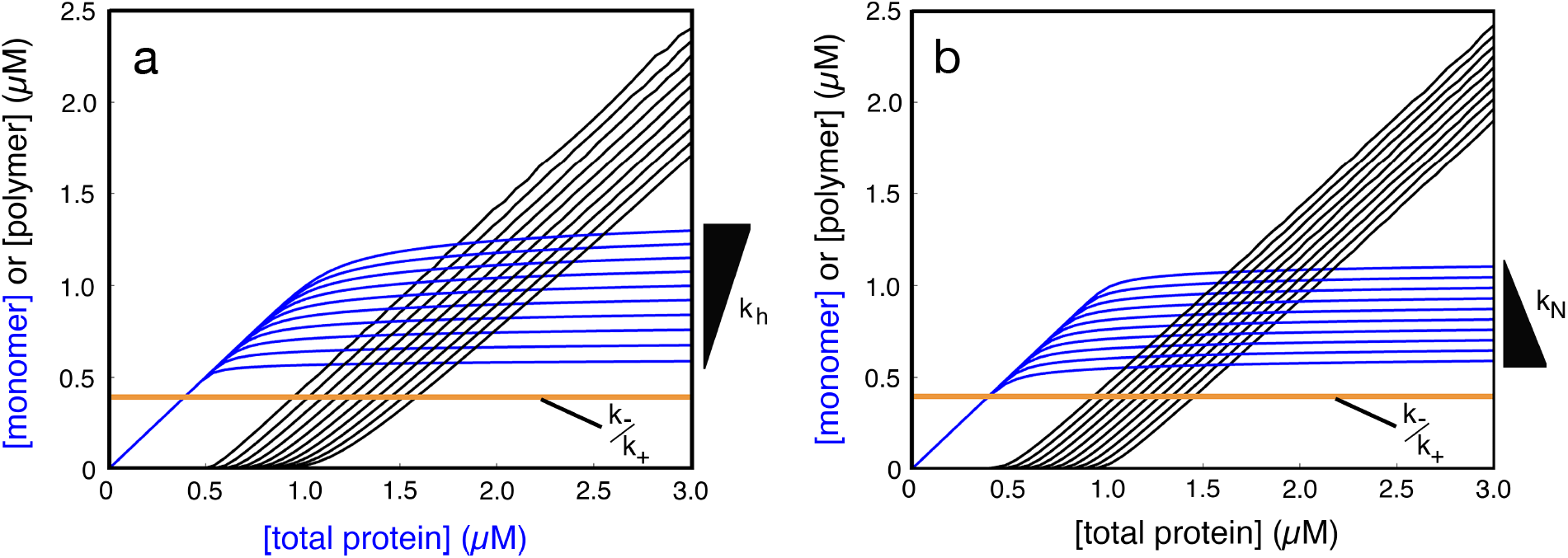
The effect of changing ATP-hydrolysis and nucleation rate constants (*k*_*h*_ and *k*_*N*_) on cc_N_, as calculated from **Eqn. 5**. (a) Increasing the ATP-hydrolysis rate constant increases the steady state monomer/critical concentration. Parameters are given in Table 1, except for *k*_*h*_ which increases, with values: 0.2, 0.3, 0.4, 0.5, 0.6, 0.7, 0.8, 0.9, 1.0, and 1.1 sec_-1_. The effect of these increasing values on the steady state monomer concentration is indicated by the black wedge. (b) Increasing the nucleation rate constant decreases the steady state monomer/critical concentration. Parameters are given in Table 1, except for *k*_*N*_ which increases, with values of: 10^-9^, 10^-8^, 10^-7^, 10^-6^, 10^-5^, 10^-4^, 10^-3^, 10^-2^, 10^-1^, 10^0^, µM^-2^sec^-1^. The effect of these increasing values on the steady state monomer concentration is indicated by the black wedge. In a and b the cc_E_ calculated from *k*_*-*_/*k*_*+*_ is marked by a yellow line.

Specifically, the cc_N_ increases by more than a factor of two as *k*_*h*_ increases from 0.2 sec^-1^ to 1.0 sec^-1^ (**Figure 5a**), while *k*_*N*_ must change by nine orders of magnitude —from 1 to 10^-9^ µM^-2^sec^-1^ — to produce a similar effect (**Figure 5b**).

Note that all sets of parameters used in **Figure 5** produce values of cc_N_ significantly higher than the critical concentration for elongation (cc_E_).

### Other models for catastrophe

The ATP-/GTP-cap model for catastrophe used in this analysis requires a single ATP-/GTP-bound monomer to stabilize the end of a filament. This is the simplest version of a class of nucleotide cap models in which hydrolysis events within a critical distance of the filament end (say, within *b* protomers) trigger disassembly (Antal, 2007a; Antal, 2007b; Dieterle, 2022). Under these conditions the critical centration for net assembly becomes proportional to the product of hydrolysis rate and critical cap size (∼*bk*_*h*_). These models can be used to fit measured values of rate and critical constants to determine the critical cap size necessary to stabilize a given filament. This is the subject of a separate paper.

### Formal similarities to the Dogterom and Leibler model

The spontaneous nucleation and ATP-/GTP-cap model in my analysis appears on the surface to be quite different from the fixed nucleation sites and concentration-dependent rescue assumed by Dogterom and Leibler (1993), but there is an important formal similarity. The nucleotide cap model assumes that ATP/GTP hydrolysis on a terminal subunit initiates rapid depolymerization. If depolymerization then uncovers additional ATP-bound protomers, the filament stabilizes and begins growing again.

These transient depolymerization events (avalanches) are small —2-5 subunits— and have the effect of modestly slowing the elongation rate (Antal, 2007a, 2007b). Global (or major) catastrophes occur only when insufficient ATP-bound protomers remain to arrest the progress of avalanches. Similar to catastrophes in the Dogterom model, avalanches occur at a fixed rate, proportional to the rate of ATP hydrolysis. Moreover, the arrest of avalanches depends on the number of ATP-bound protomers at the end of the filament, which increases with filament elongation rate —i.e. monomer concentration. This resembles a highly localized version of the concentration-dependent rescues of Dogterom, The critical transition in my spontaneous nucleation and cap model occurs when the elongation rate is high enough to create an ATP cap that prevents almost all of the avalanches from turning into major catastrophes. This is, therefore, somewhat analogous to the critical concentration in the Dogterom model, which represents the point where rescue is frequent enough to prevent filament extinction and permit unbounded filament growth. A key difference between the models is that Dogterom’s rescues are not spatially limited, but may occur anywhere along the filament. This has the effect of making filaments more stable as they grow longer. Arrests of avalanches in the cap model, in contrast are limited to the end of the filament and their frequency does not change with length or age of the filament. This partly accounts for the fact that growth in this model never becomes truly unbounded. Global catastrophe rates may drop to *near* zero but will never quite *reach* zero.

### Implications for regulation of bacterial actin-like proteins

For dynamically unstable proteins that do not interact with disassembly, recycling, and/or sequestering factors, the steady-state monomer concentration in the cytoplasm will be determined by the critical concentration for net assembly (cc_N_). If the total protein in the cytoplasm is greater than cc_N_, free filament elongation will be buffered to a constant rate: *k*_*+*_(cc_N_ - cc_E_). This appears to be true for the actin-like protein ParM. Although the cellular concentration of ParM is estimated at ∼10 µM, the elongation rate measured for filaments in bacterial cytoplasm is consistent with a free monomer concentration of 2.3 µM, the cc_N_ for ParM.

The stronger dependence of cc_N_ on catastrophe rate (∼*k*_*h*_) than nucleation rate (∼*ln*[*k*_*N*_]) means that cells can more easily change the steady-state monomer concentration by changing the catastrophe rate than by altering nucleation (Petek, 2017). This is also consistent with previous studies, which have reported suppression of catastrophe rates by filament tip-binding factors (Garner, 2007) and anti-parallel bundling (Gaythri, 2012; Petek, 2017).

## Acknowledgments

This work was supported by grants from the National Institutes of Health (1R35 GM118119) and the Howard Hughes Medical Institute. I am grateful to members of the Mullins Lab for helpful discussions and critical readings of the manuscript, especially Sam Lord, Natalie Petek, and Kristen Skruber. I am deeply indebted to Tim Mitchison for his critical reading (*r*_*crit*_) of early drafts of this manuscript and for comments which significantly improved both argument and the presentation.

